# Chemotype diversity of foliar volatile emissions in Cork oak suggests independent geographic variations in quality and quantity

**DOI:** 10.1101/2025.02.04.636386

**Authors:** Michael Staudt, Coralie Rivet, Meltem Erdogan

## Abstract

Knowledge of the intraspecific variability of volatiles produced by plants is central for estimating their fluxes from ecosystems and for understanding their evolution in an ecological and phylogenetic context. Past studies suggested that leaf volatile emissions from Cork oak (*Quercus suber* L.) exhibit a particular high degree of qualitative and quantitative polymorphism. However, the extent of inherent emission variability across its range is not known.

We investigated leaf emissions and ecophysiological variables of 241 Cork oak seedlings from ten provenances. To minimize environmental influences, emissions were determined at 30 °C and saturating light on seed-grown saplings of similar age grown under the same conditions. All individuals, except for three apparent non-emitters, released the same five monoterpenes at a rate of 2559 ± 120 ng m^-2^ s^-1^. Northern provenances tended to have higher mean emission rates and lower photosynthetic rates than southern populations, resulting in significant differences in their apparent carbon losses by volatile emissions. Independently, the emission composition varied discontinuously among individuals according to three distinct chemotypes, indicating inherent differences in the activity of two types of monoterpene synthases: one producing α-, β-pinene and sabinene, and the other limonene (plus myrcene as a by-product). Chemotype frequencies differed among provenances, particularly between South-Eastern Mediterranean and South-Western Atlantic provenances. Regarding ecophysiological leaf traits, we found no significant difference between chemotypes.

The study confirms that Cork oak is a strong monoterpene emitter, showing independent intraspecific variability in emission quality and quantity, with non-emitters being rare. Comparison of the emission variability with those reported for other oak species suggests that an ancestral Pinene/sabine chemotype has diversified within the oak subgenus *Cerris* during its radiation. This diversification is less pronounced in Cork oak than in other sympatric oaks, possibly due to differential fragmentation and expansion of their ranges in the past.

## INTRODUCTION

A large number of metabolites produced by plants can evaporate under ambient temperatures thus diffusing out of the plant organ into the air (Kesselmeier and Staudt, 1999). Ecologists and environmental chemists have studied the emissions of these volatile organic compounds (VOCs) for decades mainly for two reasons. First, the emitted VOCs can induce reactions in other receiving organisms thereby ensuring trophic and reproductive interactions potentially providing important benefits to the emitting species (e.g., Sharma *et al*., 2017; Sasidharan *et al*., 2023). Second, plant VOC emissions are the largest source of reactive carbon in the troposphere. Its air-chemical breakdown shapes the oxidative capacity and radiative properties of the atmosphere with impacts on air pollution and radiative forcing components of the climate system (Fuentes *et al*., 2000; Glasius and Goldstein, 2016). To assess their impact on air quality and climate, large-scale estimates of VOC emissions are needed (so-called emission inventories). All of them base on VOC emission capacities assumed for plant species or plant functional types (Monson *et al*., 2012). Though some plant VOCs such as methanol are likely universally produced by all plants, others such as volatile isoprenoïdes, phenyl-propanoïdes or sulphuric compounds can be highly clade specific (Kesselmeier and Staudt, 1999; Vivaldo *et al*., 2017). For example the hemiterpene isoprene, which has been identified as the strongest contributor to global plant VOC emissions, is produced at higher rates inside the leaves of 10 to 40 % of the floral species depending on the study and investigated ecosystem (Fineschi *et al*., 2013; Sharkey and Monson, 2017). Among the strongest isoprene emitters are the broadleaf trees willows, poplars and oaks. With about 500 species, the genus *Quercus* (oaks) is one of the most diversified genus within the Fagaceae family (Kremer and Hipp, 2020). Oaks are widespread all over the Northern hemisphere and can constitute the dominant species in temperate, subtropical and tropical forest ecosystems. According to the latest oak phylogeny based on genetic markers, the oak genus is subdivided into two main clades, the subgenus *Quercus* and the subgenus *Cerris* (Denk *et al*., 2017; Hipp *et al*., 2020). The subgenus *Quercus* comprises five sections containing essentially but not exclusively New World oaks (sections *Protobalanus*, *Lobatae*, *Ponticae*, *Virentes*, *Quercus*) and the subgenus *Cerris* three sections (*Cyclobalanopsis, Ilex, Cerris)* that are all old world oaks. The oldest oak ancestors probably emerged after the Cretaceous–Paleogene extinction event (Barron *et al*., 2017, and references therein) and diverged into the main clades during the Eocene, Oligocene epochs. To our knowledge, all oaks of the subgenus *Quercus* analysed for their VOC emissions have been classified as isoprene emitters due to the large dominance of isoprene in their VOC emission profiles (Supplementary information Fig. S1). Instead, oaks of the subgenus *Cerris* are predominately either strong constitutive monoterpene (MT) emitters or non-emitters. Furthermore, MT emitters typically produce a mixture of several isomers, the composition of which may vary depending on the species. Several previous studies have analysed the inter-specific variability of VOC emissions in oaks in view of its relation to oak phylogeny and to the evolution and adaptive value of VOC production (Loreto *et al*., 1998; Csiky and Seufert, 1999; Harley *et al*., 1999; Loreto, 2002; Welter *et al*., 2012; Monson *et al*., 2013). For example, the deciduous North African endemic oak *Q. afares* was suspected to origin from ancient hybridization between *Q. canariensis* and *Q. suber* based on a genetic study by Mir *et al*. (2006). Since *Q. canariensis* belongs to the subgenus *Quercus* while *Q. suber* to *Cerris*, Welter *et al*. (2012) hypothesized that different emitter types or dual isoprene MT emitters might occur in *Q. afares* populations as observed in early generation hybrids between other monoterpene and isoprene emitting oaks (Schnitzler *et al*., 2004; Staudt *et al*., 2004). The emission characteristics they found in a population of *Q. afares* reassembled that of oaks of the section *Cerris* but provided no evidence for a remnant isoprene production. Hence unlike hypothesized, these results did not support the presumed hybrid origin of *Q. afares*, which was later independently confirmed by genetic studies (Simeone *et al*., 2018; Denk *et al*., 2023).

However, the VOC emissions reported in literature for oak species and its associated assignments to an emitter type are not always consistent. For example, the Japanese oak *Q. glauca* has been described as a monoterpene emitting evergreen oak by Mochizuki *et al*. (2020), whereas as a deciduous isoprene emitter by Loreto (2002). One possible reason for inconsistency in the emission characterization of oaks is that the species identification is not always straightforward due to the occurrence of polymorphisms associated with species migrations, repeated range fragmentations and genetic introgressions following isolated hybridization events (e.g. Simeone *et al*., 2018; Leroy *et al*., 2019). In addition, isoprenoid emissions are highly plastic that strongly vary over daytime and season regulated by edaphic-climatic and ontogenetic factors (e.g., Bertin *et al*., 1997; Llusià *et al*., 2013; Ndah *et al*., 2022). Finally, as for morphological traits, intra-specific chemical polymorphism can exist leading to chemotypes differing inherently in the amount and/or the compositional fingerprint of VOCs, as evidenced in numerous studies for conifers and aromatic plants (e.g., D’Antuono *et al*., 2000; Kännaste *et al*., 2018). One oak known for exhibiting potentially both qualitative and quantitative chemotypes is the Mediterranean Cork oak (*Quercus suber* L., section *Cerris*). This emblematic evergreen oak is native to southwest Europe and northwest Africa thriving on non-calcareous substrates such as crystalline schists, gneiss, granite and sands (Toumi and Lumaret, 2001; Magri *et al*., 2007). Mature Cork oaks can be easily and unambiguously identified by their thick cork bark. Yet, earlier works conducted on a few individuals in Italy classified Cork oak as non-emitter (Steinbrecher *et al*., 1997; Delfine *et al*., 2000) while studies in Spain and Portugal as strong monoterpene emitter (Pio *et al*., 1993; 2005). In a first study investigating the intraspecific variability in two populations in France, Staudt *et al*. (2004) reported that the great majority of Cork oaks are indeed MT emitters but inherently differ in their emission profiles, i.e. chemotypes exist. In a follow-up study, Loreto *et al*. (2009) investigated the emission variability among Cork oak saplings from five provenances and confirmed the existence of these chemotypes. These authors reported divergent emission profiles between populations in continental Italy and Portugal and hypothesized that these could serve as markers of Cork oak provenances. However, the frequency of chemotypes and the exact compositional and quantitative variation of emissions within each Cork oak population was not reported in that study.

To gain more insight into the intraspecific variability of VOC emissions of Cork oak and its geographic distribution we screened the VOC emissions of Cork oak saplings originating from ten provenances in Southwest Europe and Northwest Africa. VOC emission rates were quantitatively measured at defined temperature and light conditions known as standard conditions in environmental and atmospheric sciences (Niinemets *et al*., 2010). The resulting emission rate is referred to as emission factor or basal emission rate (BER) and is integrated in models to predict VOC fluxes from land covers. Knowing how BER varies quantitatively and qualitatively within and between oak species will help to improve these emission inventories (Guenther, 2013). It may also help to identify low emitting phenotypes, ecotypes and provenances that could be used in programs of reforestation and urban greening (Sampaio *et al*., 2016; Morcillo *et al*., 2020; Chang *et al*., 2024). In order to minimize environmental influences on emissions we used saplings of similar age that were grown together and kept under the same conditions during the experimental period. By focusing on the innate determinants of chemical polymorphism in VOC emissions, we aimed to answer the following questions:

Do inherent qualitative or quantitative chemotypes exist in Cork oak populations as indicated by previous studies?

What is the distribution of the putative chemotypes in the different Cork oak provenances?

Do Cork oak chemotypes or provenances differ in ecophysiological properties?

How chemodiversity in Cork oak compares to that of other related monoterpene emitting oak species and their phylogeny?

## MATERIALS AND METHODS

### Plant culture

The Cork oak acorns were collected from different trees in ten sites representing the following provenances (approximate geographic latitude, longitude coordinates): **Akfadou** Mountains in Northern Algeria (36.66, 4.62); **Catalonia** in the French Western and Spanish Northern Mediterranean (42.47, 2.92), **Corsica** island (France) around Porto Vecchio (41.63, 9.22); **Espadan** mountains in Eastern Central Spain (39.87, -0.30); **Kroumerie** Mountains in Northern Tunisia (36.79, 8.69); **Maâmora** forest region at the Northern Atlantic coast of Marocco (34.07, -6.67); **Maremma** coastal region in western central Italy north of Grosseto (42.92, 11.15); **Sardinia** island (Italy) around Iglesias (39.33, 8.53), **Setubal** district in South-west Portugal (38.27, -8.31); **Var** department in the French Eastern Mediterranean (43.25, 6.42).

Except of the Akfadou population (see below), all plants were grown and kept in plastic pots with a mixture of argilo-calcareous soil, sand and peat moss compost (pH = 5,5-6,5). Potted plants were kept outside in the institute garden, where they were irrigated during summer and occasionally fertilized. The screening experiment was run from middle of July to middle of October. To reduce eventual weather and seasonal effects on emissions, saplings were transferred beginning of July in air-conditioned and light controlled greenhouse. Target day/night temperature was 25/20 °C and the minimum light level during lighted hours ca 500 μmol m^-2^ s^-1^ Photosynthetic Photon Flux Density (PPFD). Saplings were randomly selected for emission screening, ensuring, however, to alternate among provenances. To assess the plasticity of VOC emissions, we made on five randomly chosen individuals 4 replicate measurements during the experimental period (ca. every 3 weeks). In addition to the saplings grown in pots, we assessed the chemical profiles of 18 mature Cork oak trees of the Akfadou Mountains planted in our institute garden (see Welter *et al*., 2012). To do so, we cut branches from the trees, whose cut end were immediately immerged in a bucket with water and recut under water to remove eventual xylem embolism. The bucket with the branch was then brought to the lab and assayed for VOC emission as described below for the potted saplings.

### VOC emission measurements

Foliar VOC emissions along with CO_2_/H_2_O gas exchanges was measured with a dynamic environmentally controlled exposure system consisting of two small flat chambers (Vol. 105 mL) run with an air flow rate of 0.7 L min^-1^ (Staudt and Lhoutellier, 2011). The mature terminal leaves of a twig of a sapling were mounted in a chamber and adapted to chamber conditions for at least 30 min. Chamber air temperature was set to 30 °C and incident PPFD to ca. 1500 μmol m^-2^ s^-1^. Chamber air was delivered by an compressor (Ingersoll Rand, model 49810187), which was cleaned and dried by a clean air generator (Airmopure, Chromatotec, St Antoine, France) and a charcoal filter, and then re-humidified by directing an adjustable portion of the air stream through a bypasses with washing bottles. The average air to leaf vapour pressure deficit during measurements was 0.026 ± 0.005 mol mol^-1^. CO_2_ and H_2_O air concentrations of the chamber airs was monitored using a LiCOR 640 infra-red gas analyzer at a flow rate of 0.2 L min^-1^.VOCs were sampled by directing a portion (0.1 L min^-1^) of the chamber airs for 10 min into Perkin Elmer adsorbent cartridges containing ca. 290 mg Tenax TA. Immediately after VOC sampling, the LiCOR 640 analyzer was switched to the chamber incoming air and the enclosed leaves were collected to determine their projected area (Delta-T Areameter MK2, Delta-T Devices Ltd) and dry weight after having dried them for 3 days at 60 °C (microbalance Mettler PM200, Mettler-Toledo SA). Empty chamber measurements were regularly conducted to detect eventual background VOCs.

Cartridges were analyzed with a Varian CP3800/Saturn2000 gas chromatograph mass spectrometer (GC-MS) equipped with a Perkin Elmer Turbomatrix thermodesorption unit. Trapped VOCs were thermally desorbed (15 min at 230 °C) on a cold trap (-30 °C) containing a small amount of Tenax TA, and subsequently thermally injected into a Chrompack Sil 8 CB low bleed capillary column (30 m × 0.25 mm × 0.25 μm). The GC oven temperature program was 3 min at 40 °C, 3 °C min^-1^ to 100 °C, 2.7 °C min^-1^ to 140 °C, 2.4 °C min^-1^ to 180 °C, 6 °C min^-1^ to 250 °C. Carrier gas was helium (1.0 ml min^-1^). In addition to the offline VOC measurements, the chambers were connected to an online gas chromatograph (AirmoVOC C2-C6 Gas Chromatograph, Chromatotec, St Antoine, France) for the detection of isoprene (see Staudt and Visnadi (2023), for a more detailed description).

### Data treatment and statistics

The VOC emission rate was calculated as the difference between the VOC air concentration in the chamber enclosing leaves and the mean concentration measured in the empty chambers multiplied by chamber airflow rate, and divided by the projected leaf area or by the leaf dry mass. The calculation of the leaf CO_2_/H_2_O gas exchange variables net-photosynthesis (A), transpiration (E), water vapour conductance (G_H2O_), internal CO_2_ concentration (Ci) was made according to von Caemmerer and Farquhar (1981) using the CO_2_ and H_2_O mixing ratios of the air leaving and entering the chambers. We further deduced from these data the water use efficiency (WUE = A E^-1^) and the intrinsic water use efficiency (iWUE = A G_H2O_^-1^). Variation in iWUE and Ci at given [CO_2_], light and temperature indicates adjustments in the limitations of the leaf’s efficiency to fix atmospheric CO_2_ other than stomatal opening (e.g. Rubisco activity, mesophyll conductance), whereby iWUE decreases and Ci increases with decreasing efficiency. Finally, we calculated the ratio between VOC emission and A as the percent mole of the actually assimilated carbon lost by emission (C-loss). The formation of constitutive monoterpenes and isoprene inside chloroplasts is directly linked to photosynthetic processes delivering the basic carbon substrates and energetic cofactors for the chloroplast isoprenoid biosynthesis pathway. Labelling studies with ^13^CO_2_ have shown that the major fraction if not all of their carbon comes from the ongoing CO_2_ fixation in the Calvin-Benson-Bassham Cycle (e.g., Loreto *et al*., 1996; for review see e.g. Lantz *et al*., 2019). Thus, the C-loss reflects the actual percent allocation of photosynthates to volatile isoprenoid production. Given this link, we discarded all measurements from the data set, in which A was very low (ca. < 1 µmol m^-2^ s^-1^) in order to avoid identifying false non-emitters.

We analysed relative emission data (percentage composition) with hierarchical cluster analysis and factorial discriminant analysis to differentiate chemotypes. Correspondence analyses were applied to the contingency table of the frequency of chemotypes in the ten provenances to test for differences in their distributions. Subsequently, we applied a Mantel test to test the correlation between the dissimilarity in the chemotype frequency among provenances and their geographical distances. To test whether provenances and chemotypes differed in ecophysiological variables we first run MANOVAs followed by single factor ANOVAs or Kruskal Wallis tests in case data did not fill the necessary requirements. Differences between groups were considered to be significant at the level α = 0.05. Posthoc Bonferroni and Dunn tests were used for pairwise comparison of categories with adjusted significance levels (α) of 0.0011 and 0.0083 for provenance and chemotype, respectively. Pearson correlation analyses and scatter plots were performed to test the covariation among variables. All statistical analyses were done with addinsoft (2021) XLSTAT statistical and data analysis solution.

Unless otherwise stated, the degree of data scattering (variances) of mean values are reported as ± standard error (SE = standard deviation divided by the square root of the number of replicates) and as relative variation coefficient (VC = standard deviation divided by the mean x 100).

## RESULTS

Of 241 individuals, only three were found to be non- or very low VOC emitters. The average total VOC emission rate with these non-emitters included was 2655 ± 121 (VC: 71 %) ng m^-2^ s^-1^ or 55.8 ± 2.4 µg g^-1^ h^-1^ (VC: 68 %) and the apparent photosynthetic carbon investment for VOC emissions (C-loss) 2.3 ± 0.2 % (VC: 110 %). Excluding the three non-emitters (n = 238), the mean total emission rate was 2698 ± 122 (VC: 70%) ng m^-2^ s^-1^ or 56.5 ± 2.4 µg g^-1^ h^-1^ (VC: 66 %), and the C-loss was 2.4 ± 0.2 % (VC: 109 %). The major emitted VOCs were α-pinene, sabinene, β-pinene, myrcene und limonene. These five MTs were detected in all emitting trees and accounted on average 95% of the total emission (2559 ± 120 ng m^-2^ s^-1^; 53.7 ± 2.4 µg g^-1^ h^-1^; n = 238). In addition, the monoterpenes, α-thujene, α-terpinene, 1,8-cineol, (Z)*-* and (E)*-β*-ocimene, the sesquiterpenes germacrene D and *β*-caryophyllene, and the green leaf volatiles (Z)-3-hexenol and (Z)-3-hexenyl acetate were irregularly detected, mostly at trace amounts except ocimenes, which sometimes was released at high rates (>500 ng m^-2^ s^-1^). Isoprene, if any, was only occasionally detected at trace levels (<1 ng m^-2^ s^-1^). Given the irregular appearance of ocimenes and minor VOCs, we focus in the following on the five major MTs.

In all ten populations, their percentage composition varied among individuals expressing a discontinuous variation pattern (Fig. 1). The emissions of the majority (52.5 %) was mainly composed of α-, β-pinene and sabinene, whereas the profile of 13 % contained mainly limonene. The remainder (34.5 %) emitted all four compounds in higher proportions. Repeated measurements conducted on five trees showed that the individual emission profiles were quite stable even though the overall quantity varied considerably among plant replicates (Supplementary Information Fig. S2). This suggest that the compositional fingerprint is largely inherent, whereas the actual production rate is more subject to plasticity even under environmentally controlled conditions. Hierarchical cluster analysis grouped the emission profiles into three clearly distinguished chemotypes as confirmed by factorial discriminant analysis (Supplementary Information Fig. S3). These chemotypes are hereafter referred to as Pinene/Sabinene type, Limonene type and Mixed type, and the three individuals lacking these emissions as Non-emitter. Correlation analyses of the relative emissions of single compounds shows that α, β-pinene and sabinene scaled all positively to each other (*R*: 0.85 to 0.89) and all negatively to limonene (*R*: -0.94 to -0.97) (see Supplementary Information Fig. S4). Myrcene proportions were weakly positively correlated to limonene (*R*: 0.55) and weakly negatively to pinenes and sabinene (*R*: -0.52 to -0.68).

**Fig. 1.**
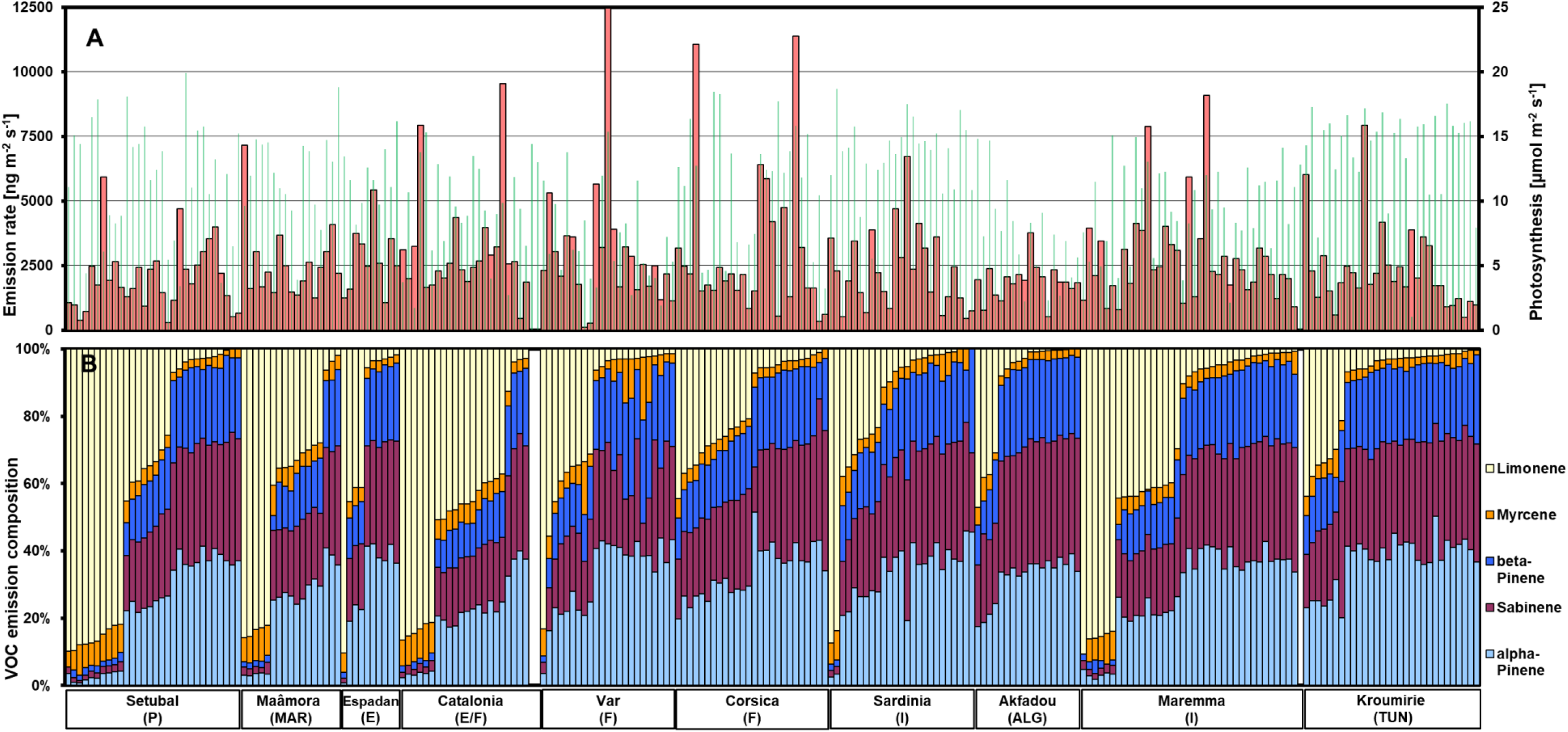
Leaf monoterpene emissions from 241 cork oak saplings originating from ten provenances. (A) Emission rate of the sum of the five main compounds (red columns) and photosynthesis rate (green columns) measured at 30°C and ca. 1500 incident PPFD; (B) relative proportions of the five main monoterpenes. The data were first sorted according to the origin of the trees (provenance) and then according to the proportion of limonene in their emissions. Note that three seedlings emitted no VOCs or only traces, although their leaves were fully photosynthetically active.

Based on Chi-square statistic, correspondence analyses suggest real differences between the populations in terms of their chemotype frequencies (*P* < 0.001). The most diverged emission profiles were between the Eastern provenances Akfadou, Kroumirie, Corsica and the Western provenances Maâmora, Catalonia and Setubal, the former group lacking completely the Limonene type, which was most present in the latter group (Fig. 2). The profile of the Maremma population with the highest number of screened individuals was the closest to the mean profile of all. Results of a Mantel test suggest that the dissimilarity in the chemotype frequency among provenances is not correlated with their geographical distances (*P* = 0.053, *R*^2^ = 0.1).

**Fig. 2.**
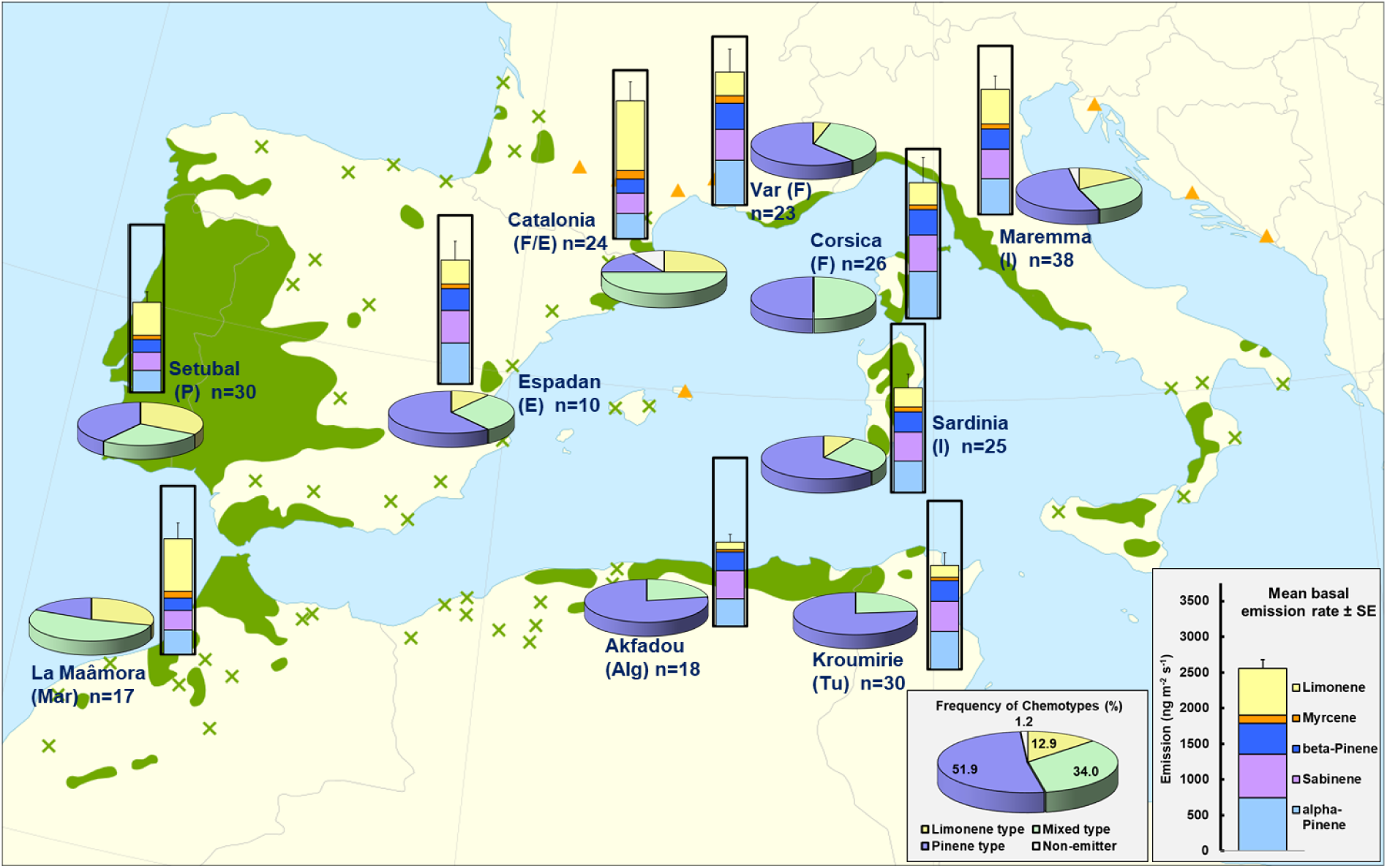
Overview of the mean emission rate (column charts) and frequency of chemotypes (pie charts) of the ten provenances within the Cork oak distribution range (n = number of plant replicates). In all column plots the Y-axes are at the same scale and the error bars show the SE for the sum of the emissions of the five major monoterpenes. The big insert charts in grey show the means across all investigated provenances. The distribution map is taken from Caudullo *et al*. (2017). The green colored areas in the map are the continuous native Cork oak ranges and the green crosses isolated native populations. Orange triangles indicate introduced and naturalized Cork oak populations.

MANOVAs applied on the leaf gas-exchange variables (A, E, G_H2O_, Ci, WUE, iWUE), SLW, BERs and associated C-losses suggest significant differences between populations (*P* < 0.0001) but no significant difference between the three major chemotypes (*P* = 0.117). Single factor ANOVAs or equivalent Kruskal Wallis tests confirmed the absence of significant differences among chemotypes for any variable except of the extremely low mean emission rate and C-loss of the Non-emitter type compared to the emitting chemotypes (Table 1). In contrast, single factor tests revealed significant population differences for the variables A (*P* < 0.0001), E (*P* < 0.0001), G_H2O_ (*P* < 0.0001), iWUE (*P* = 0.0001), Ci (*P* = 0.002) and SLW (*P* < 0.0001). The provenance differences in A were essentially to differences in G_H2O_ though not exclusively as indicated by provenance differences in iWUE. Provenances differed also in BER (*P* = 0.032 for the sum of major MTs) but overall in the associated photosynthetic carbon investments (C-loss, *P* < 0.0001). The lowest mean C-losses were observed in the populations from Kroumerie, Setubal and Sardinia and the highest values in the northern provenances Var, Maremma and Catalonia (non-emitters excluded). The trees of the Algerian provenance (Akfadou) had the lowest mean emission rate but had also low mean photosynthesis rate. Since measured on cut branches, its comparison with other populations should be taken with caution. The provenance mean BERs per leaf surface correlated negatively with their mean Ci (*R* = -0.83, *P* = 0.003) and moderately positively with their mean iWUE (*R* = 0.67, *P* = 0.034) (Fig. 3). However, using mean BERs per leaf dry weight instead of per leaf area reduced correlations both to Ci (*R* = - 0.61, *P* = 0.060) and iWUE (*R* = 0.51, *P* = 0.129). In fact the provenance means of Ci tended to be negatively correlated with SLW (*R* = -0.61, *P* = 0.059). Besides, there was no significant correlation between the mean BERs and the means of SLW, A, T, G_H2O_ or WUE (*P* > 0.2; Supplementary Information Table S1).

**Fig. 3.**
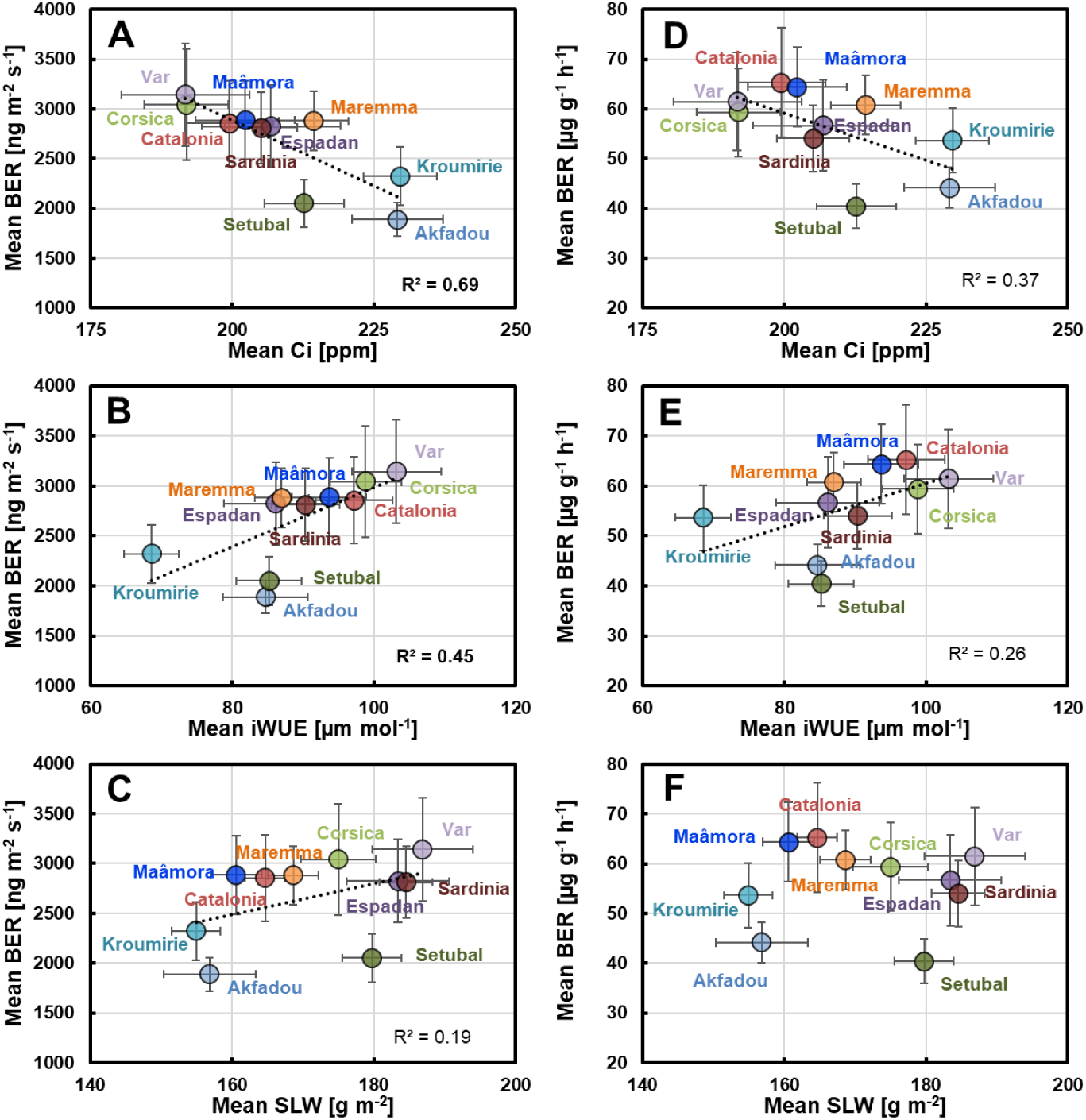
Plot of mean basal monoterpene emission rates (BER) per provenance against mean leaf internal CO_2_-concentrations Ci (A, D), mean intrinsic water use efficiencies iWUE (B, E) and mean specific leaf weights SLW (C, F). The left panels (A, B, C) show the relationships to the BERs per leaf area, the right panels (D, E, F) to the BERs per leaf dry mass. Dotted lines with coefficients of determination R^2^ show results from linear correlations. Correlation coefficients (R) and corresponding P-values are summarised in the Supplementary Information Table S1.

**Table 1.** Basal emission rates and CO_2_/H_2_O gas exchange variables of *Q. suber* saplings from 10 provenances. Values are means ± SE per provenance (upper table) and per Chemotype (lower table). The line below each sub-table summarizes the P-values of ANOVA (A) or Kruskal-Wallis (K) tests for differences between Provenance or Chemotypes (**Significant** at P < 0.05). Superscript letters denote which of the categories are different based on pairwise comparisons using the Bonferroni test (A) and Dunn test (K). Note that these post-hoc tests apply adjusted significance levels lower than 0.05 (for Chemotype α = 0.0083, for Population α = 0.0011). Therefore, the differences between pairs of categories are sometimes reported as insignificant even if the global test result is significant at α = 0.05.

|  | Major MT<br>emission | Total MT<br>emission | A | G<sub>H2O</sub> | SLW | Ci | WUE | iWUE | C-loss<br>(major MTs) |
| --- | --- | --- | --- | --- | --- | --- | --- | --- | --- |
| Provenance (n) | [ng m<sup>-2</sup> s<sup>-1</sup>] | [ng m<sup>-2</sup> s<sup>-1</sup>] | [μmol m<sup>-2</sup> s<sup>-1</sup>] | [mmol m<sup>-2</sup> s<sup>-1</sup>] | [g m<sup>-2</sup>] | [mmol m<sup>-2</sup> s<sup>-1</sup>] | [mmol mol<sup>-1</sup>] | [μmol mol<sup>-1</sup>] | [%] |
| Sardinia (25) | 2320 ± 311<sup>a</sup> | 2812 ± 359<sup>a</sup> | 13.6 ± 0.6<sup>c</sup> | 158 ± 9<sup>cd</sup> | 184 ± 4<sup>a</sup> | 205 ± 6<sup>ab</sup> | 3.47 ± 0.14 | 90 ± 5<sup>ab</sup> | 1.33 ± 0.21<sup>a</sup> |
| Setubal (30) | 2012 ± 236<sup>a</sup> | 2051 ± 242<sup>a</sup> | 11.7 ± 0.8<sup>bc</sup> | 144 ± 11<sup>bc</sup> | 180 ± 4<sup>abc</sup> | 213 ± 7<sup>ab</sup> | 3.25 ± 0.14 | 85 ± 5<sup>ab</sup> | 1.70 ± 0.38<sup>a</sup> |
| Maâmora (17) | 2572 ± 349<sup>a</sup> | 2886 ± 393<sup>a</sup> | 12.0 ± 0.8<sup>bc</sup> | 129 ± 7<sup>bc</sup> | 161 ± 4<sup>bcd</sup> | 202 ± 9<sup>ab</sup> | 3.58 ± 0.20 | 94 ± 5<sup>ab</sup> | 1.71 ± 0.28<sup>ab</sup> |
| Espadan (10) | 2752 ± 418<sup>a</sup> | 2825 ± 415<sup>a</sup> | 11.8 ± 0.7<sup>abc</sup> | 156 ± 28<sup>bc</sup> | 183 ± 7<sup>ab</sup> | 207 ± 12<sup>ab</sup> | 3.34 ± 0.22 | 86 ± 7<sup>ab</sup> | 1.84 ± 0.33<sup>ab</sup> |
| Kroumirie (30) | 2309 ± 289<sup>a</sup> | 2322 ± 291<sup>a</sup> | 13.9 ± 0.7<sup>c</sup> | 210 ± 9<sup>d</sup> | 155 ± 4<sup>d</sup> | 229 ± 7<sup>b</sup> | 3.37 ± 0.15 | 67 ± 4<sup>a</sup> | 2.03 ± 0.87<sup>a</sup> |
| Corsica (26) | 3001 ± 560<sup>a</sup> | 3043 ± 559<sup>a</sup> | 11.0 ± 0.9<sup>abc</sup> | 116 ± 11<sup>abc</sup> | 175 ± 5<sup>abcd</sup> | 192 ± 7<sup>a</sup> | 3.71 ± 0.17 | 99 ± 5<sup>b</sup> | 2.11 ± 0.29<sup>ab</sup> |
| Akfadou (18)<sup>2</sup> | 1876 ± 166<sup>a</sup> | 1889 ± 167<sup>a</sup> | 7.4 ± 0.9<sup>ab</sup> | 99 ± 16<sup>ab</sup> | 157 ± 6<sup>cd</sup> | 229 ± 8<sup>ab</sup> | 4.21 ± 0.42 | 85 ± 6<sup>ab</sup> | 2.32 ± 0.35<sup>ab</sup> |
| Maremma (37) | 2783 ± 295<sup>a</sup> | 2957 ± 293<sup>a</sup> | 9.4 ± 0.5<sup>ab</sup> | 114 ± 8<sup>ab</sup> | 169 ± 4<sup>abcd</sup> | 214 ± 6<sup>ab</sup> | 3.45 ± 0.15 | 87 ± 4<sup>ab</sup> | 2.40 ± 0.25<sup>b</sup> |
| Catalonia (24) | 3065 ± 431<sup>a</sup> | 3112 ± 430<sup>a</sup> | 9.4 ± 0.8<sup>ab</sup> | 100 ± 9<sup>ab</sup> | 165 ± 3<sup>abcd</sup> | 200 ± 8<sup>ab</sup> | 3.59 ± 0.19 | 97 ± 6<sup>b</sup> | 3.31 ± 0.66<sup>b</sup> |
| Var (23) | 2957 ± 518<sup>a</sup> | 3142 ± 516<sup>a</sup> | 7.2 ± 0.8<sup>a</sup> | 71 ± 7<sup>a</sup> | 187 ± 7<sup>a</sup> | 192 ± 11<sup>a</sup> | 3.58 ± 0.20 | 103 ± 6<sup>b</sup> | 3.52 ± 0.58<sup>b</sup> |
| **P<sub>Provenance</sub> (Test)** | **0.032 (K)<sup>1</sup>** | 0.151 (K)<sup>1</sup> | < **0.0001 (K)** | < **0.0001 (K)** | < **0.0001 (K)** | **0.002 (K)** | 0.361 (K) | **0.0001 (K)** | < **0.0001 (K)** |
| **Chemotype (n)** |  |  |  |  |  |  |  |  |  |
| Limonene (31) | 2503 ± 314<sup>a</sup> | 2545 ± 322<sup>a</sup> | 10.8 ± 0.8 | 126 ± 13 | 170 ± 4 | 208 ± 7 | 3.39 ± 0.16 | 93 ± 5 | 2.47 ± 0.52<sup>a</sup> |
| Mixed (82) | 2562 ± 197<sup>a</sup> | 2666 ± 195<sup>a</sup> | 10.7 ± 0.4 | 124 ± 6 | 169 ± 3 | 207 ± 4 | 3.58 ± 0.10 | 92 ± 3 | 2.08 ± 0.16<sup>a</sup> |
| Pinene 125 | 2580 ± 173<sup>a</sup> | 2767 ± 179<sup>a</sup> | 10.8 ± 0.4 | 136 ± 6 | 172 ± 2 | 211 ± 4 | 3.49 ± 0.09 | 86 ± 2 | 2.51 ± 0.27<sup>a</sup> |
| None (3) | 3 ± 1<sup>b</sup> | 28 ± 14<sup>b</sup> | 13.4 ± 0.5 | 139 ± 6 | 177 ± 4 | 191 ± 21 | 4.32 ± 0.35 | 97 ± 8 | 0.02 ± 0.01<sup>b</sup> |
| **P<sub>Chemotype</sub> (Test)** | **0.028 (K)<sup>1</sup>** | **0.023 (K)<sup>1</sup>** | 0.705 (K) | 0.570 (K) | 0.822 (A) | 0.822 (A) | 0.153 (K) | 0.235 (A) | **0.025 (K)** |
| **Total 241** | **2527 ± 120** | **2665 ± 123** | **10.8 ± 0.3** | **131 ± 4** | **171 ± 2** | **209 ± 3** | **3.53 ± 0.06** | **89 ± 2** | **2.21 ± 0.16** |

<sup>1</sup> Tests were run with emission rates expressed per leaf surface. Using emission rates expressed per leaf dry mass reduce the significance level of provenance effects for Major MT emissions ( $P = 0.092$ ) and increase it for Total MT emissions ( $P = \mathbf{0.032}$ ). It does not change the significance level for differences between chemotypes for Major MT emissions ( $P = \mathbf{0.027}$ ) and increases it for Total MT emissions ( $P = \mathbf{0.024}$ ).
<sup>2</sup> Measurements were made on cut branches collected from adult trees, which may have affected CO<sub>2</sub>/H<sub>2</sub>O gas exchanges and VOC emissions.

## DISCUSSION

### Quantity - variability in BER

The amount of MTs produced inside the chloroplasts of oak leaves depends on the amount of active MT synthases and the availability of its unique substrate geranyl-diphosphate produced from photosynthetic triose-phosphate and energetic cofactors in the MEP pathway (Loreto *et al*., 1996, Fischbach *et al*., 2002; Niinemets *et al*., 2002; Staudt *et al*., 2022). With a mean BER of ca. 50 µg C g^-1^ h^-1^ (20 nmol m^-2^ s^-1^) consuming more than 2% of the net-assimilated carbon, our results place Cork oak among the highest MT emitters in the Mediterranean area, rivalling the BERs known for strong isoprene emitters and exceeding those reported for sympatric conifers and aromatic shrubs (Owen and Hewitt, 2001; Bracho-Nunez *et al*. 2013). Nevertheless, few individuals did not release volatile isoprenoids in relevant amounts. Non-emitters among strong MT emitting individuals have already observed in populations of Cork oak (Staudt *et al*., 2004), as well as in populations of *Q. afares* (Welter *et al*., 2012) and recently of *Q. phylliraeoides* (Chang *et al*., 2024).

To minimize the phenotypic plasticity in MT production we measured and adapted plants to the same assay conditions using in great majority saplings of same age that were grown together. Despite of this, the BER of the 238 emitting saplings that all released the same five principle MTs varied by up to two orders of magnitude (VC ca 70%). Repeated measurements conducted on same individuals over the same period revealed only up to five time differences in BER (mean VC ca 40%) indicating that a significant portion but not all of the total between-plant variability in BER was inherent. Correlation analyses and comparison of variances provide some clues to the underlying reasons. The BER variability was somewhat lower when expressed on leaf dry weight (CV 66.3%) than on projected leaf area (CV 70.1%) indicating that leaves having a higher SLW tended to have higher per leaf area VOC emissions. Population differences in SLW were also associated with population differences in BER, Ci and iWUE. Several mechanisms operating at different scales can explain these associations. Typically, high SLW leaves are better acclimatized to high irradiance (sun leaves), resist more to harsh environmental conditions and have a longer lifespan (Niinemets 2007, Poorter *et al*. 2009). Their anatomy often displays an increased per area volume of chloroplasts, in which photosynthesis and monoterpene biosynthesis take place. As a result, the per leaf area BER and A are increased but not necessarily its ratio, i.e. the C-loss. In addition to anatomical differences, high SLW leaves can show an increased activity of rate limiting photosynthetic enzymes to fix more efficiently CO_2_ at high radiation. This would lead to lower Ci and higher iWUE. As shown for several isoprene and monoterpene emitting trees species, low Ci can stimulate the formation and emission of isoprenoids inside the chloroplasts by enhancing the biosynthesis of their precursors in the MEP pathway (Niinemets *et al*., 2021; Staudt *et al*., 2022; Sahu *et al*., 2023; and other references therein). Finally, the level of active MT synthases inside chloroplasts might inherently vary among individuals unrelated to Ci and photosynthetic processes. In both cases, the apparent C-loss is expected to change too. Indeed in our study, the provenance means of Ci scaled negatively with mean SLW and BERs (Figure 3, Supplementary Information Table S1). However, it explained less than half of the BER variability suggesting that a mix of metabolic and morpho-anatomical adjustments were shaping the mean BER of populations to different extents. For example differences in Ci could be the major cause for the much lower mean BER of Setubal compared to Corsica, which otherwise were similar (SLW, A; see Table 1). On the contrary, it does not explain the higher mean BER of Maremma saplings, which had similar mean Ci and lower mean SLW compared to Setubal. Thus, the very low BER of this Portuguese population might be attributed to low MT-synthase activity plus reduced allocation of photosynthates in MT synthesis (high Ci) leading to an overall low mean C-loss. Low mean C-losses were also observed in the populations from Sardinia, Estaban, Maâmora and Kroumerie that in addition to more moderate BER tended to have higher mean A compared to the Northern provenances Maremma, Var and Catalonia. This might indicate a north-south up-regulation of resource investment that optimizes carbon gain under high light and temperature conditions at the expense of MT production (Rasulov *et al*., 2014; Monson *et al*. 2020).

### Quality - variability in the emission composition

Beside and independent of the variability in the emitted quantities, the emissions of the five major emitted compounds exhibited discrete compositional variations classifying them into three non-plastic chemotypes, wherein three of the five MTs clearly co-varied. Many MT synthases are multiproduct enzymes synthesizing several MT isomers in defined proportions (Degenhardt *et al*., 2009). Thus, the recurrent discontinuous emission pattern observed in the Cork oak populations could be due to differences in the activity of only two terpene synthases, one producing predominantly α-, β-pinene and sabinene, and one producing predominantly limonene. Myrcene is probably a by-product of both enzymes, but especially of the limonene-producing enzyme. The activity of each putative MT synthase contributing to the final emission blend of the three chemotypes would depend on genomic, proteomic but most likely on transcriptomic differences as commonly observed in other studies (Padovan *et al*., 2017 and references therein). Co-regulation of the actual MT production rates by substrate availability (see above) can explain the absence of the significant differences among the mean BERs of the three emitting chemotypes and the independence of population differences in BER and chemotype frequency. Similar Pinene/sabinene and Limonene chemotypes were already described in previous studies on Cork oak (Staudt *et al*., 2004; Loreto *et al*., 2009), as well as on *Q. ilex* (Holm oak; Staudt *et al*., 1999, 2004), *Q. afares* (African oak; Welter *et al*., 2012) and *Q. coccifera* (Kermes oak; Staudt *et al*., 2022). Chemotypes contrasting in their limonene and pinene proportions were also reported for distantly related species such as *Abies grandis* (Katoh and Croteau, 1998), *Cannabis sativa* (Fischedick *et al*., 2010) and Cotton (Clancy *et al*., 2023). Concerning Mediterranean oaks, several studies indicated a west-east gradient in the frequency of chemotypes, with the Limonene type being more prevalent in western populations (Staudt *et al*., 2004, Loreto *et al*., 2009, Staudt *et al*., 2023; and other references therein). Consistent with this, most genetic studies reported a west-east differentiation pattern in their gene pools with two or more major clusters depending on the marker (Simeone *et al*., 2018; Pina-Martins et al, 2019; Sousa *et al*., 2022, 2023; and other references therein). In the present study, some western provenances showed indeed a high frequency of the limonene and mixed type (Setubal, Maâmora, Catalonia), which in turn was rare or absent in the two eastern North African populations, characterised by a relatively homogeneous composition dominated by the Pinene/sabinene type. However, the Limonene type was also found in some eastern populations, including the most northeastern Maremma provenance, which is not consistent with the use of the Pinene/sabinene chemotype as a marker for continental Italian provenances proposed by Loreto *et al*. (2009). Thus, our results show that the distribution of chemotype within the Cork oak range is more complex than a matter of longitudinal gradient or geographical distance.

A major limitation in comparing emission studies with phylogenetic studies is that the latter base on neutral genetic markers, frequently chloroplast DNA markers that are only maternally inherited (Holderegger *et al*., 2006). MT synthase genes are nuclear encoded, hence maternally and parentally inherited, and their expression to form volatiles may not be neutral but affects the plants ability to withstand environmental stresses (e.g., Weraduwage *et al*., 2024). Nevertheless, several results of our study are worthy to discuss in an evolutionary context. One question that arises here is to what extent the common diversification pattern in the emissions of MT-emitting oaks results from a common ancestral polymorphism, which then barely evolved during the radiation of the species, or whether it is the result of a recent parallel evolution of a highly versatile trait. Indeed MT production as other groups of much diversified “secondary” plant metabolites can evolve rapidly often involving gene duplication followed by neofunctionalization owing to few mutations that changed the shape of the enzymes’ active site (for reviews, see e.g. Tholl, 2006; Weng *et al*., 2012; Christianson, 2017). Comparing the MT emissions reported in the present and in our previous studies for *Q. suber*, *Q. ilex*, *Q. coccifera* and *Q. afares* reveals that the proportions among the three positively correlated MTs (α-, β-pinene and sabinene) produced by putative pinene/sabinene synthases differ between the oaks of the section *Ilex* (*Q. ilex* and *Q. coccifera*), and the oaks of the section *Cerris* (*Q. suber* and *Q. afares*) but not or less so between the oaks of a same section.

As can be seen in Fig. 4, the two *Ilex* oaks exhibit consistently a higher ratio in the β-pinene and sabinene proportions than the *Cerris* oaks. Furthermore, the emissions of the two *Ilex* oaks express other chemotypes in addition to the Pinene/Sabinene, Limonene, Mixed types, namely a Myrcene type and a 1,8-Cineol chemotype, both being most frequent in Kermes oak. In the Kermes oak study (Staudt *et al*.; 2023), the analytical separation of pinene stereoisomers (enantiomers) provided evidence that pinenes with sabinene are formed by at least two active MT synthases. Regarding the literature about the emission composition of other MT emitting oaks of the subgenus *Cerris*, Csiky and Seufert (1999) mentioned sabinene, α-pinene and β-pinene being the main emitted compounds of the semi-deciduous East-Mediterranean oak *Q. ithaburensis* (section *Cerris*). Among East Asian oaks, Mochizuki *et al*. (2020) and Chang *et al*. (2024) reported that the major compounds released by *Q. glauca*, *Q. myrsinifolia* (both section *Cyclobalanopsis*) and *Q. phillyraeoides* (section *Ilex*) are α-pinene followed by sabinene and β-pinene in proportions apparently similar to those we observed in the Pinene/sabinene type. Mochizuki *et al*. (2020) also mentioned a very similar emission composition for *Castanopsis sieboldii* and *Castanea crenata*, two species of other *Fagaceae* genera. These observations may indicate that pinene/sabinene emission is an ancestral trait within the subgenus *Cerris*, which was modified *a posteriori* during subgenus expansions, whereby MT production in Holm and Kermes oak became more diversified compared to Cork oak and possibly other oaks of the section *Cerris*. According to Denk *et al*. (2023), their common ancestors origin in northern East Asia and date to the early Eocene epoch under temperate climates. Subsequently the sections split off during migrations. The *Cerris* section expanded northwestern over Siberia, whereas oaks of the section *Ilex* south-westward in subtropical climates of southern China and south-eastern Tibet, from where they moved further west along the pre-Himalayan mountains. *Ilex* and *Cerris* oaks joined later in western Eurasia along with some isoprene emitting oaks from eastern North America of the section *Quercus* (e.g. *Q. robur*, *Q. petrea*, *Q. pubescens*). Several species adapted congruently to the advent of summer dry climate around the evolving Mediterranean Basin, during which their VOC emission characteristics might have been modified. Loreto *et al*. (2009) hypothesized that the west-east distribution of limonene and pinene chemotypes in Cork oak could coincide with an early genetic divergence of Cork oak populations starting 25 Mill years ago by tectonic microplate movements as suggested by Magri *et al*. (2007). The interpretation by Magri and co-workers was however challenged by later genetic studies, which related its intraspecific genetic diversity to differential introgressive hybridizations with other sympatric oak species (essentially with *Q. cerris* in the east and with *Q. ilex* in the west) and/or to repeated range contractions during the Pleistocene glaciation cycles (Lumaret and Jabbour-Zahab, 2009; Simeone *et al*., 2018, Pina-Martins *et al*., 2019; López de Heredia, 2020). A common result of genetic studies is the lower genetic diversity of Cork oak compared to other sympatric oaks (Sousa *et al*., 2023; and other references therein), which is consistent with the reduced chemical polymorphism we observed in its VOC emissions. Based on nuclear alloenzyme variation, Toumi and Lumaret (1998) observed a higher genetic diversity in the western Iberian populations compared to several south-eastern provenances including from North Africa. Accordingly, we observed the less diversified emission profiles in the Algerian and Tunisian populations. With a relative narrow ecological amplitude, Cork oak has a more restricted and fragmented range than *Q. ilex*, *Q. coccifera* and other sympatric oaks such as *Q. cerris*, which resist better to freezing and snow charge, grow well on limestone and have more continuous and larger eastward extended ranges in the Mediterranean basin (Toumi and Lumaret, 2001; Caudillo *et al*. 2017). Therefore, local bottlenecks and enhanced genetic drifts during repeated retractions and isolations of populations in ice age refugia along with human selection have particularly reduced the genetic diversity in Cork oak. We hypothesize that this genetic erosion is also reflected in its impoverished emission profile compared to *Q. ilex* and *Q. coccifera*. Introgressive hybridization events might have been contributed to qualitative and quantitative emission diversifications. Simeone *et al*. (2018) inferred on the Italian peninsula ingressive forms of the non-emitting oak *Q. cerris* into *Q. suber*, possibly explaining the occurrence of non-emitting Cork oak individuals observed in continental Italy (Steinbrecher *et al*., 1997). Regarding the introgression with Holm oak, seven individuals of the Pinene/sabinene chemotype in our study emitted β-pinene in higher proportions than sabinene plus relatively large proportions of myrcene, as is typical for the emissions of the two *Ilex* oaks (Fig. 4B). Interestingly, all but one of these saplings were from the Var provenance.

**Fig. 4.**
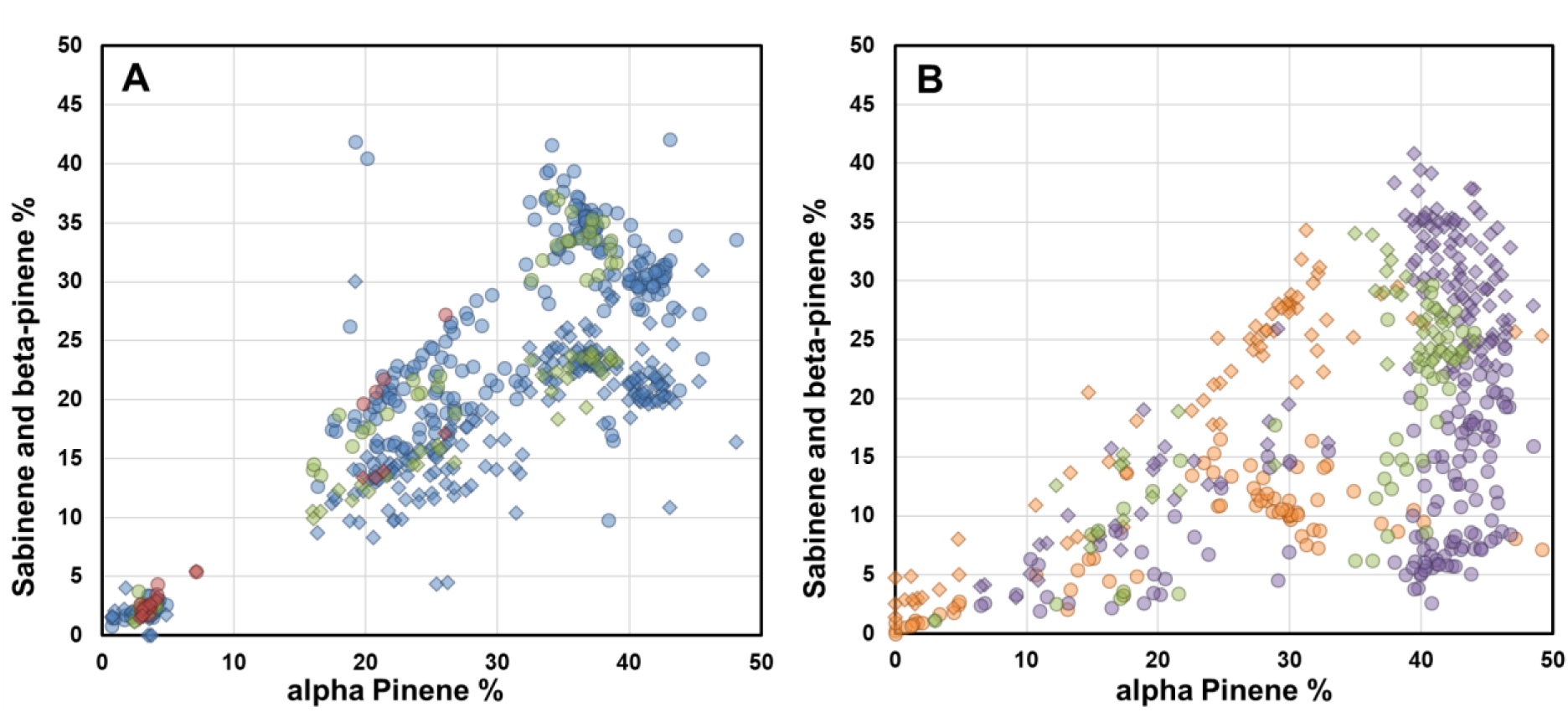
Plots of the proportions of sabinene (circles) and b-pinene (diamonds) against the proportions of a-pinene in the emissions of A: *Q. suber* (Cork oak) and *Q. afares* (African oak) and in the emissions of B: *Q. ilex* (Holm oak) and *Q. coccifera* (Kermes oak). The shown emission data were taken from different studies. In A: blue: *Q. suber* (this study ); green: *Q. suber* (Staudt *et al*., 2004): brown: *Q. afares* (Welter *et al*., 2012). In B: purple: *Q. ilex* (Staudt *et al*., 1999); green: *Q. ilex* (Staudt *et al*., 2004); red: *Q. coccifera* (Staudt and Visnadi, 2023). In all the emissions, Pinene/Sabinene, Limonene and Mixed chemotypes were observed. However, in the emissions from *Q. suber* and *Q. afares*, the proportions of sabinene are higher than those of b-pinene, whereas in *Q. ilex* and *Q. coccifera*, the opposite is observed.

## CONCLUSIONS AND OUTLOOK

Our study confirms that Cork oak is a strong MT emitter, but it also shows strong non-plastic inherent intraspecific variability in BER, including the rare occurrence of individuals that do not emit. BER varied considerably even within provenance populations, such that the difference between mean BERs of provenances was barely significant. Nevertheless, there were clear differences in the apparent allocation of photosynthetic carbon to MT production, which was lower in the southern provenances. Correlations with ecophysiological variables such as Ci and SLW indicate that both anatomical and metabolic adaptations are involved. Beside BER variability, VOC emissions of plants displayed a discontinuous variation in their composition separating them in three distinct chemotypes. These chemotypes were unevenly distributed across the range, with limonene-producing chemotypes more common in some western populations and Pinene/sabinene chemotype more common in south-eastern populations. Comparison with the chemical diversity of emissions reported for other oak species suggests that the Pinene/sabinene chemotype is an ancestral trait common to the subgenus *Cerris*, which was modified during clade radiations, whereby Cork oak emissions became less diversified than the emissions of its congeners, possibly due to strong recurrent fragmentation and restrictions of its range and interspecific genetic exchanges. Regarding leaf ecophysiological traits, we found no significant difference between chemotypes.

Together, our results offer new perspectives for understanding the extent and origins of VOC emission diversity in oaks as well as for the correct assignment of BERs in emission models, especially for models coupled to photosynthetic processes (Morfopoulos *et al*., 2013). Furthermore, the non-emitting individuals we found could provide an opportunity to reduce regional VOC pollution by propagating them in reforestation programs, although their rarity suggests that they are subject to negative selection. To better understand the underlying mechanisms it would be interesting to determine the activity of monoterpene synthases, measure other metabolites of the isoprenoid pathway in Cork oak chemotypes and conduct genome comparisons for terpene synthases across different oak species. Half-sib populations of non-emitters and specific chemotypes could be generated to elucidate the inheritance mode of MT emissions, and be used for testing their ecological and phenological characteristics such as growth rate or vulnerability to abiotic and biotic stresses (e.g., Sampaio *et al*., 2016, Bertic *et al*., 2023, Sanchez-Osorio *et al*., 2024).

## Supporting information

Staudt etal Preprint Supplementary Information

## SUPPLEMENTARY INFORMATION

**Figure S1.** Overview on the interspecific variability of isoprenoid emissions within the genus *Quercus*.

**Figure S2.** VOC emission rates and composition and photosynthesis rates from four replicate measurements carried out on five Cork oak seedlings during the experimental period.

**Figure S3.** Dendrogram from hierarchical cluster analysis and observation diagram from factorial discriminant analysis showing that the variation in the composition of the five major monoterpenes emitted by 238 Cork oak individuals can be categorised into three different chemotypes.

**Figure S4.** Plot of the proportions of α-pinene versus the proportions of the four other main MTs in the emissions from 238 Cork oak saplings.

**Table S1.** Matrix with results of Pearson correlation among the provenance mean values of the basal emission rate, specific leaf weight, leaf internal CO_2_ concentration, photosynthesis, transpiration, water vapour conductance, water use efficiency and intrinsic water use efficiency.

## ACKNOWLEDGEMENTS

The authors thank B. Buatois, P. Daubin, D. Degueldre and E. Hilbig, for their valuable assistance in plant cultivation, setting up and running experiments. We further wish to thank all colleagues who helped us to find or collected Cork oak acorns of different provenances, namely R. Piazetta, Drs M. Abourouh, H. Almeida, J. Aronson, R. Joffre, R. Lumaret, H. Merouani and D. Mousain.

