## Supplementary material for "Chemotype diversity of foliar volatile emissions in Cork oak suggests independent geographic variations in quality and quantity": Staudt etal Preprint Supplementary Information

CEFE, CNRS, EPHE, IRD, Univ Montpellier, Montpellier, France

**Figure S1** (see next page). Interspecific variability of isoprenoid emissions within the genus ***Quercus***. The shown oak phylogeny was adapted from Denk et al. (2017), after which the genus is subdivided into the two subspecies (***Cerris***, ***Quercus***) comprising respectively three (*Cyclobalanopsis*, *Ilex*, *Cerris*) and five sections (*Lobatae*, *Protobalanus*, *Ponticae*, *Virentes*, *Quercus*). The approximate number of species within each section is given in brackets. For the sections that have only a few species (*Cerris*, *Protobalanus*, *Ponticae*, *Virente*), all species have been listed, including those for which we could not find information on their constitutive VOC emissions. VOC emission data were compiled considering the following publications (and other references therein):

Bertin et al. 1997; doi: 10.1016/S1352-2310(97)00080-0
Steinbrecher et al., 1997, doi: 10.1016/S1352-2310(97)00076-9)
Loreto et al., 1998; doi: 10.1007/s004420050520
Csiky and Seufert, 1999; doi: 10.1890/1051-0761(1999)009[1138:TEOMOA]2.0.CO;2
Kesselmeier and Staudt, 1999; doi: 10.1023/A:1006127516791
Harley et al., 1999; doi: 10.1007/s004420050709
Guenther et al., 1999; 10.1016/S1464-1909(99)00062-3
Loreto, 2002; doi: 10.1078/1433-8319-00033
Lim et al., 2011; doi: 10.1016/j.atmosenv.2011.01.066
Okumura et al., 2008; doi: 10.2525/ecb.46.257
Tani and Kawawata, 2008; doi: 10.1016/j.atmosenv.2008.01.059
Tani et al., 2011; doi: 10.1016/j.atmosenv.2011.08.003
Steinbrecher et al., 2009; doi: 10.1016/j.atmosenv.2008.09.072
Welter et al., 2012; doi: 10.1093/treephys/tps069
Monson et al., 2013; doi: 10.1111/pce.12015
Yaman et al. 2015; doi: 10.4209/aaqr.2014.04.0082
Mochizuki et al., 2020; doi: 10.1080/13416979.2020.1779425
Bao et al., 2023; doi: 10.1016/j.envpol.2022.120886
Staudt et al., 2023; doi: 10.1525/elementa.2023.00043
Baek et al., 2024; doi: 10.1016/j.atmosenv.2024.120654
Tani et al., 2024; doi: 10.1186/s40645-024-00645-8
Yu et al., 2024; doi: 10.1016/j.atmosenv.2023.120238

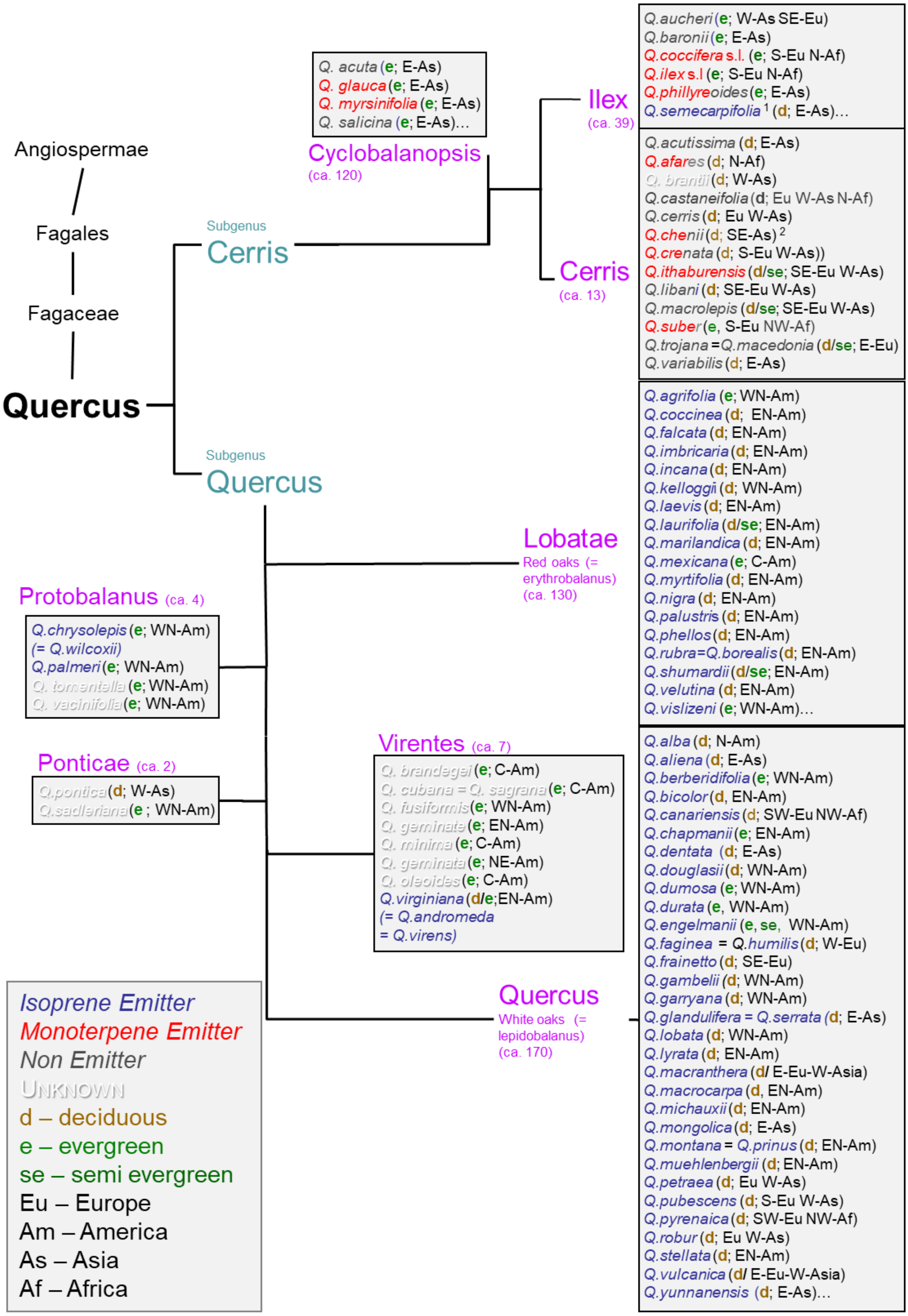


[1] based on a single study by Loreto et al. (1998), which measured VOC emissions from an unknown number of seedlings grown from acorns collected in the field.

[2] classified as a non-isoprene emitter in Harley et al. (1999), without giving details on whether this species emits monoterpenes or not.

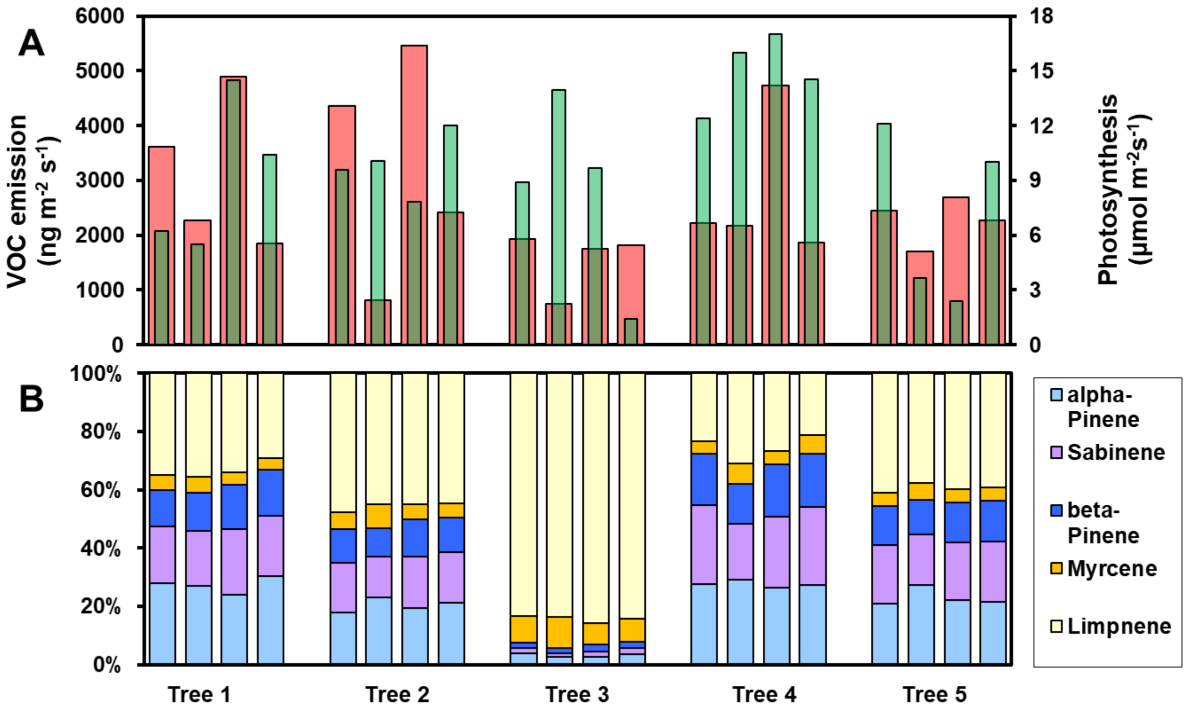


**Figure S2.** Repeated measurements of leaf VOC emissions and photosynthesis from five cork oak saplings. (A) Emission rate of the sum of the five main VOCs (red columns) and photosynthesis rate (green columns). (B) Relative proportions of the five main VOCs. During the experimental period, four replicate measurements were carried out on each sapling at 30°C and approx. 1500 incident PPFD on annual leaves of different twigs. The results show that the compositional fingerprint of the VOC production of each tree is very stable, while the quantities emitted are subject to greater fluctuations.

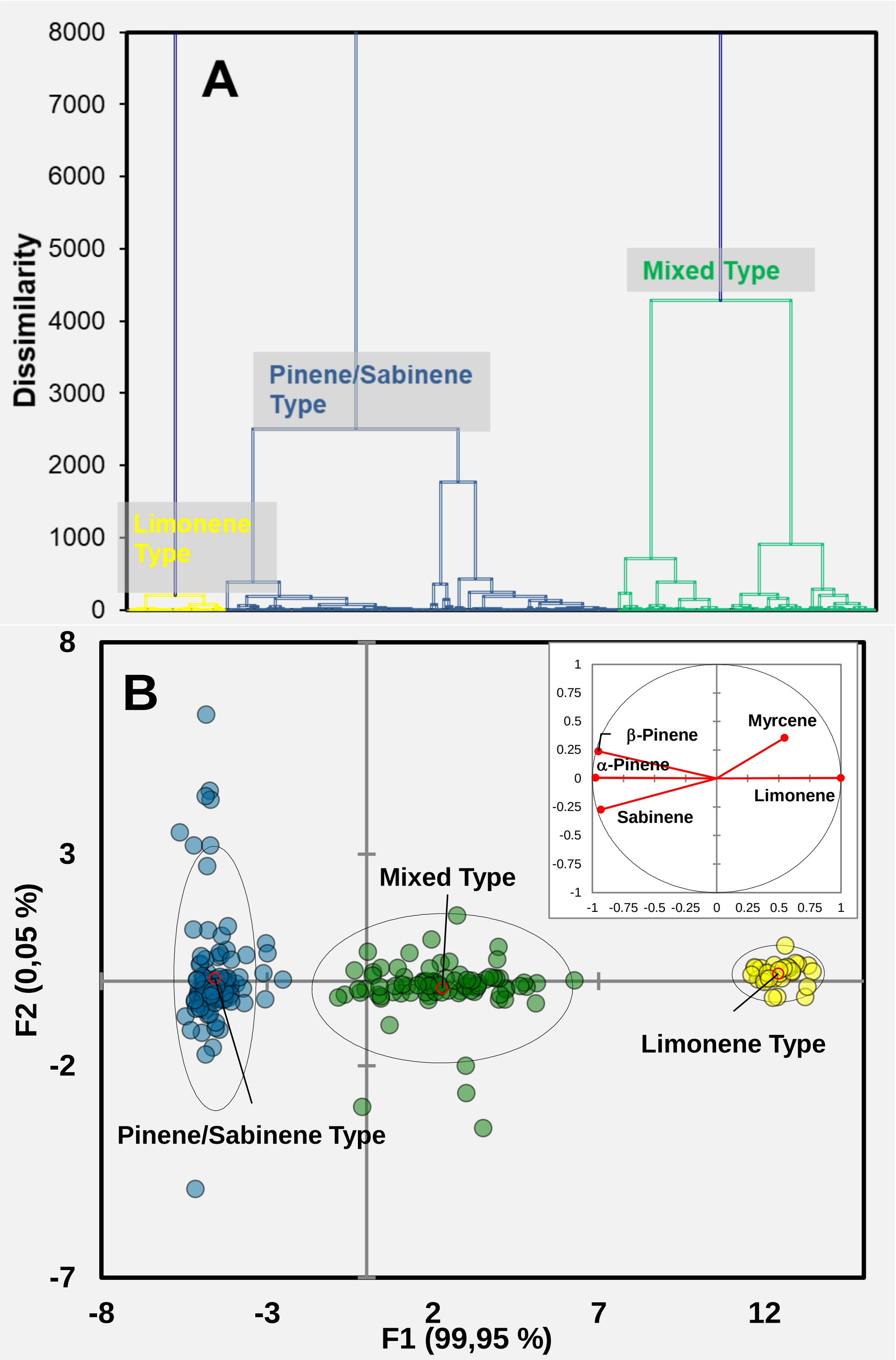


**Figure S3.** The compositional variation of the 5 major monoterpenes emitted from 238 cork oak individuals clusters in three distinct chemotypes: Limonene type (13 %, yellow), Pinene/Sabinene type (52.5 %, blue) and Mixed type (34,5 %, green). Panel A shows the dendrogram obtained from hierarchical cluster analysis using Ward's technique. Panel B shows the observation plot resulting from Factorial Discriminant Analyses (Centroids in red). More than 99 % of the observed variance is represented by Factor 1 (x-axis) correlated with the proportions of limonene, pinenes and sabinene (small inserted graph).

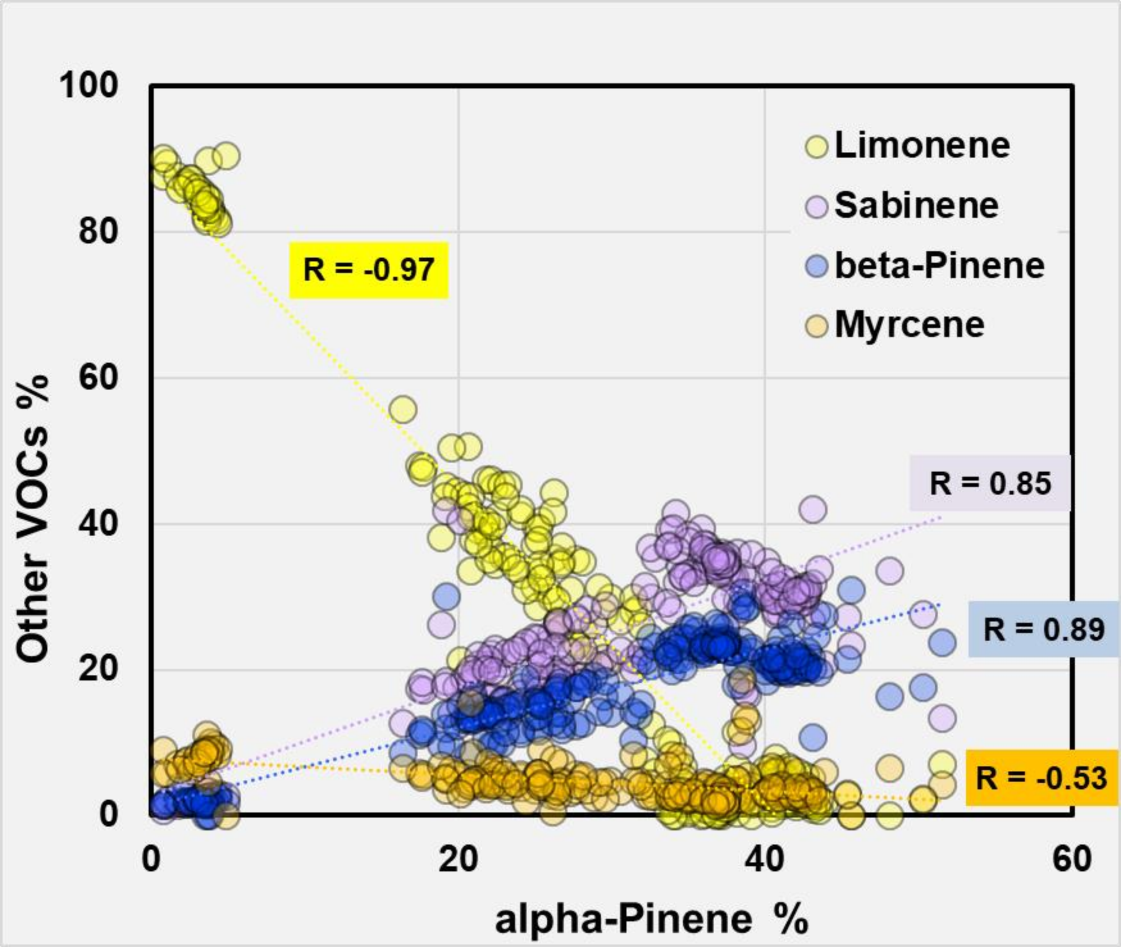


**Figure S4.** Plot of the proportions of α-pinene against the proportions of the four other major VOCs emitted from 238 Cork oak saplings (sum of 5 = 100 %). Dotted lines show best-fit correlations with correlation coefficients R assuming linear relationships. The proportions of pinenes and sabinene are strongly positively correlated in the emissions (R = 0.85 to 0.89), indicating that these three monoterpenes are always synthesised and emitted together in cork oak leaves, while limonene is apparently produced independently from pinenes and sabinene (R = -0.94 to -0.97). Mycene, which is only emitted in small amounts, could be predominantly a by-product of limonene synthesis (R = 0.55).

**Table S1.** Matrix with results of Pearson correlation among the provenance mean values of the basal emission rate (BER), specific leaf weight (SLW), leaf internal $CO_2$ concentration (Ci), photosynthesis (A), transpiration (E), water vapour conductance ($G_{H2O}$), water use efficiency (WUE) and intrinsic water use efficiency (iWUE). The upper right half of the table shows the **R-values** and the lower left half the corresponding **P-values**. Two values are given for the BERs: the first one stands for the BERs per leaf area while the second one in brackets stands for the BERs per leaf dry mass (see also Figure 3 in the main text). Values in Bold are significant at alpha = 0.05.

| **P\R** | **BER $m^{-2}$ ($g^{-1}$)** | **SLW** | **Ci** | **A** | **E** | **$G_{H2O}$** | **WUE** | **iWUE** |
|---|---|---|---|---|---|---|---|---|
| **BER $m^{-2}$ ($g^{-1}$)** | | 0.44 (-0.01) | **-0.83** (-0.61) | -0.04 (-0.08) | -0.05 (-0.12) | -0.31 (-0.28) | -0.24 (-0.10) | **0.67** (0.51) |
| **SLW** | 0.21 (0.98) | | -0.61 | 0.01 | 0.08 | -0.18 | -0.41 | 0.48 |
| **Ci** | **0.003** (0.06) | 0.059 | | 0.151 | 0.188 | 0.50 | 0.15 | **-0.90** |
| **A** | 0.91 (0.82) | 0.99 | 0.68 | | **0.976** | **0.90** | -0.61 | -0.51 |
| **E** | 0.88 (0.75) | 0.82 | 0.60 | **<0.0001** | | **0.92** | **-0.72** | -0.56 |
| **gH2O** | 0.39 (0.43) | 0.62 | 0.14 | **0.000** | **0.000** | | -0.53 | **-0.81** |
| **WUE** | 0.50 (0.78) | 0.23 | 0.67 | 0.061 | **0.020** | 0.11 | | 0.22 |
| **iWUE** | **0.034** (0.13) | 0.16 | **0.000** | 0.14 | 0.093 | **0.004** | 0.55 | |
